# Response of oriental xerophytes to the occidental industrial revolution

**DOI:** 10.1101/173013

**Authors:** Sara Manafzadeh, Yannick M. Staedler, Hamid Moazzeni, David Masson, Jürg Schönenberger, Elena Conti

## Abstract

Since the Industrial Revolution, human activities have contributed substantially to climate change, by adding CO_2_ to the atmosphere, especially since the mid-20^th^ century (the “Great Acceleration”). Climatic change does not have the same impact on different regions of the Earth, neither in the recorded past, nor in models of the future. Therefore, to anticipate on these changes, we need to understand and be able to predict the possible responses of the different regional vegetations of the world to these changes, and most significantly to increased drought conditions. The aim of this study is to understand the response of the xerophytes of the immense oriental Irano-Turanian bioregion to post-industrial global warming and to compare it with the response of the xerophytes of the neighbouring occidental Mediterranean bioregion. We measured stomata index and stomata density from 83 herbarium sheets (coll. 1821-2014) from species of the non-succulent xerophyte, *Haplophyllum*. We tested for differences before and after the “Great Acceleration” in both bioregions. SI decreased in the occidental species (significant only for abaxial leaf side), whereas SI significantly increased in the oriental species (both sides). We suggest that changes in both occidental and oriental species are linked to atmospheric CO_2_ due to the different constraints that act on their growth. In light-limited occidental species, atmospheric CO_2_ caused the stomata index decrease, whereas in the predominantly water-limited oriental species, increased drought stress and temperature (climatic change) caused stomata index increase. In conclusion, we propose that whereas atmospheric CO_2_ directly caused a decrease in stomata index in occidental xerophytes, it indirectly caused an increase in stomata index in oriental xerophytes *via* climate change (increase in aridity, drought stress, and temperature). This study highlights the considerable potential of research based on historical herbarium collections to answer ecological questions, especially regarding climatic change.

## 1. Introduction

The Earth has experienced a constantly changing climate in the time since plants first evolved. Carbon dioxide (CO_2_) is an important greenhouse gas and higher or lower atmospheric levels of CO_2_ have been linked, respectively, to warmer or cooler climates in the past (Beerling and Chaloner, 1993; Crawley and Berner, 2001; Kürschner et al., 2001; van der Burgh et al., 1993). Over the period 1860-2017, the global atmospheric partial pressure of CO_2_ increased from 280 to 407,39 ppm (figure 1a). Yearly CO_2_ release has consistently increased over time, but strongly accelerated after WWII, an event called the “Great Acceleration” (Hibbard et al., 2006; McNeill and Engelke, 2016; Steffen et al., 2015); see Fig. 1b). As a consequence, the Earth surface temperature increased and more prolonged dry periods alternate with more intensive rainfall events (Bartholomeus et al., 2012; Solomon et al., 2007), thereby affecting vegetation characteristics (Bates et al., 2008; Wegehenkel, 2009).

**Figure 1.**
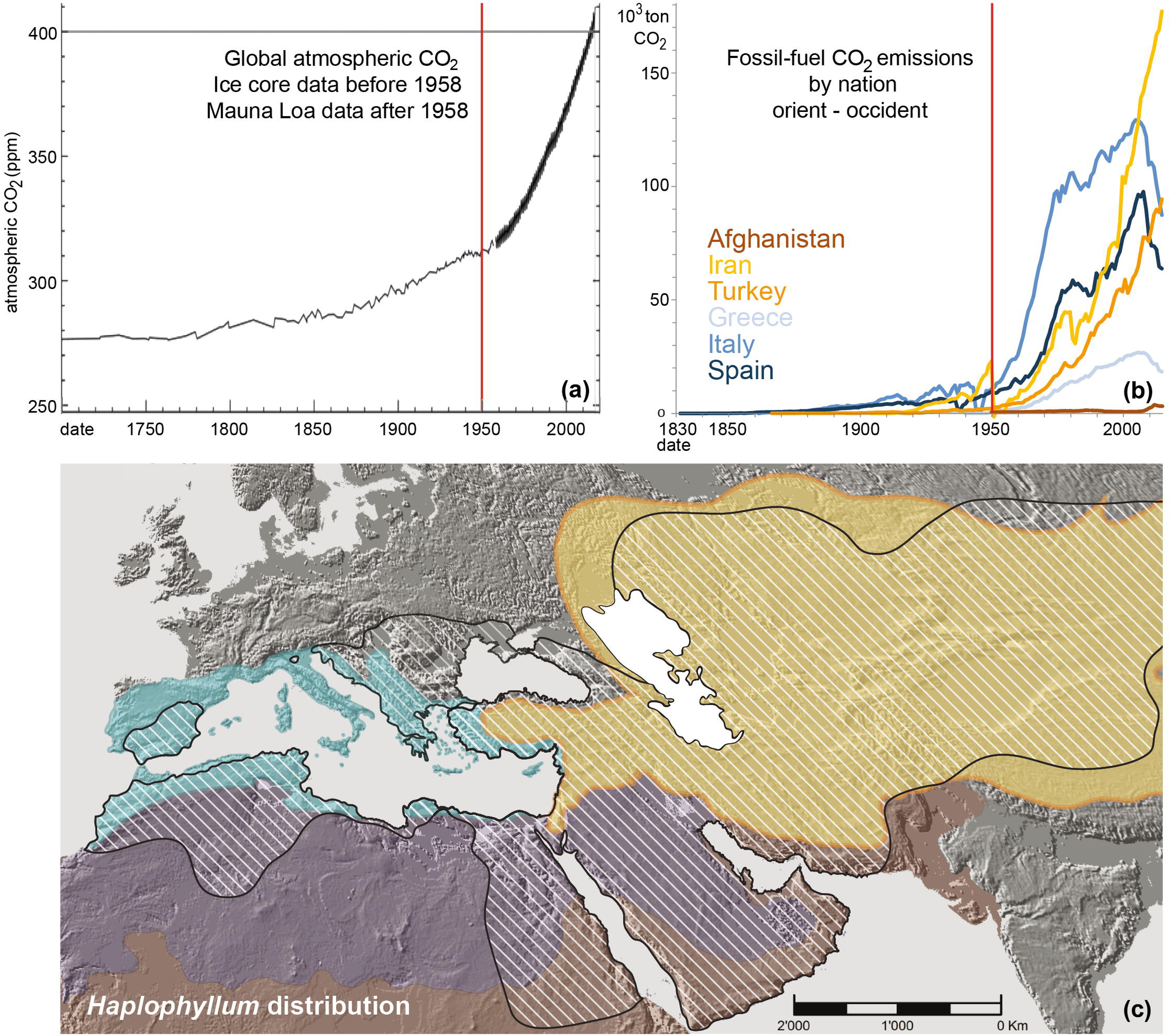
CO_2_ *versus* time and region. (a) Atmospheric CO_2_ since the XVIII^th^ century from ice core data and MaunaLoa observatory (modified from Scripps Institution of Oceanography http://bluemoon.ucsd.edu/co2_400/co2_800k_zoom.png). (b) CO_2_ emission for selected countries from Occident (inside Mediterranean floristic region) and Orient (inside Irano-Turanian floristic region; data from (Boden and Andres, 2017)). (c) *Haplophyllum* in the Orient and Occident. *Haplophyllum* distribution, hatched; IT region, yellow; Mediterranean region, blue; Saharo-Arabian region, violet; Sudano-Zambesian region, brown (modified from (Manafzadeh et al., 2014)).

Climatic change does not have the same impact on different regions of the Earth, neither in the recorded past, nor in models of the future (Pachauri et al., 2014). Biogeographic origins and constraints based on lineage may also influence the way plants respond to global change in the Anthopocene, including elevated CO_2_, warming, and changes in rainfall (Cavender-Bares et al., 2016). Therefore, to anticipate on these changes, we need to understand and be able to predict the possible responses of the different regional vegetations of the world to these changes, and most significantly to increased drought conditions. At the broadest scale, the area of the Earth can be divided into distinct bioregions, also called floristic regions in plant sciences (Kreft and Jetz, 2010; Takhtajan, 1986).

Among floristic regions, the Irano-Turanian (IT) region is one of the largest: it occupies the vast drylands of Eurasia, covering *ca.* 30% the surface of the continent, from easternmost Mongolia to the Sinai Peninsula (Manafzadeh et al., 2016); Fig. 1c). Unlike the Mediterranean region (i.e., Occident) which enjoys a mild Summer-dry climate (Takhtajan, 1986), the IT region (i.e., Orient) is characterised by high continentality, low winter temperature, and strong precipitation seasonality (Djamali et al., 2012). Given these climatic conditions, plants adapted to low-water availability, i.e., xerophytes are predominant. In contrast with the neighbouring Mediterranean floristic region where vegetation is dominated by forests and shrubland (maquis), trees are rare in the IT region, and geophytes and low shrubs are dominant (Takhtajan, 1986). Moreover, the largely industrial occidental countries (inside the Mediterranean floristic region) and the mostly developing oriental countries (inside the IT floristic region) have both shown the great acceleration in increase of CO_2_ emissions since the 1950 (figure 1b).

Despite their importance in the evolution of the Eurasian flora, the adaptation of the xerophytes of the immense IT region to post-industrial global warming has never been studied. Moreover, no study has yet investigated whether the responses of xerophytic elements to climatic change differ between the Mediterranean (occidental) and the Irano-Turanian (oriental) xerophytes.

The xerophytic genus *Haplophyllum* A. Juss. from the Citrus family (Rutaceae) is a characteristic genus of the IT floristic region (Takhtajan, 1986; Zohary, 1973) whose evolution is closely connected with the geological and climatic development of the arid and semi-arid areas of the IT region (Manafzadeh et al., 2014). *Haplophyllum*’s main centre of diversity lies in the IT region, but it is also present in the Mediterranean floristic region (figure 1c; (Salvo et al., 2011)). *Haplophyllum* comprises 68 species of perennial herbs and shrubs with elliptical to linear leaves, growing overwhelmingly in open, dry, sunny habitats, in steppe and semi-desert areas in the Orient and in open woodlands, maquis, and forests in the Occident (Salvo et al., 2011; Townsend, 1986). *Haplophyllum* is thus a perfect case study to understand the effect of post-industrial warming on Eurasian xerophytes.

Stomata are the microscopic structures on the epidermis of leaves formed by two specialized guard cells that control the exchange of water vapor and CO_2_ between plants and the atmosphere (Willmer and Fricker, 1996). According to their water storing strategies, xerophytes have either relatively low stomatal density (SD: stomata number per surface unit) in succulent xerophytes, or relatively high SD in nonsucculent xerophytes, as in *Haplophyllum* (Maximov, 1929; Volkens, 1887). CO_2_ increase has been shown to mostly cause a decrease in SD (reviewed in (Royer, 2001)). Stomata density is easy to measure and is often used when epidermal cells cannot be counted accurately; however, SD can be affected both by the initiation of stomata and the expansion of epidermal cells, which in turn is a function of many variables (e.g., light, temperature, water status, position of leaf on crown, and intra-leaf position). Carbon dioxide plays a stronger role in stomatal initiation than in epidermal cell expansion; therefore Salisbury (Salisbury, 1927) introduced the stomatal index (SI), which normalizes for the effects of this expansion. The SI is defined as the number of stomata per unit of leaf area divided by the number of epidermal cells plus number of stomata per unit leaf area (see equation below).

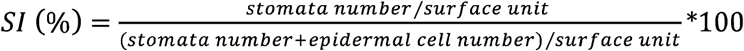

The SI is considered to be fairly constant within the leaves of a single plant and exhibits a clear response (usually an increase) to increased atmospheric CO_2_ (Beerling and Royer, 2002; Greenwood et al., 2003; Retallack, 2001; Royer, 2001; Willmer and Fricker, 1996).

Gas exchange usually takes place on the shaded underside of the leaf, the abaxial side, where stomata are more abundant and more uniformly distributed than the upper side of the leaf, the adaxial side, which can also be completely devoid of stomata. However, no significant difference in response between the two different sides of the leaves was found in experimental increases on atmospheric CO_2_ (Royer, 2001). We thus expect that post-industrial increase in CO_2_, especially after the great acceleration, should decrease SI and SD in both the occidental (Mediterranean) and oriental (IT) plants. We also expect that this CO_2_ increase leads to a stronger decrease in SI and SD in the abaxial than in the adaxial surface of the leaves. In order to assess the impact of CO_2_ increase on leaf parameters, we tested for differences between values of SI and SD on each leaf side before and after the great acceleration (the date of which we set at 1950).

We therefore studied herbarium collections of both historical and recent specimens of oriental (IT) and occidental (Mediterranean) xerophytes (i.e., *Haplophyllum*), in order to answer the following questions: (i) what is the stomata response in Eurasian xerophytes (i.e., *Haplophyllum*) to post-industrial increase of atmospheric CO_2_ concentration? and (ii) is this response dependant on floristic regions?

## 2. Material and methods

### 2.1. Taxon sampling

Two hundred and seventeen herbarium sheets from the herbarium of the Natural History Museum in Vienna (W) and from the Herbarium of the University of Zurich (Z) were sampled. Due to poor preservation state and the difficulties to remove cuticular waxes (see below) data acquisition was possible for 83 of the 217 specimens which were collected from 1821 to 2014. Our final dataset includes 16 species of *Haplophyllum,* 12 of which occur in the Orient (inside the IT floristic region) and three of which occur in the Occident (inside the Mediterranean floristic region). One species occurs in both the Orient and the Occident (table S1). The herbarium authorities did not allow us to destructively sample more than one leaf per sheet.

### 2.2. Data acquisition

The leaves were removed from the herbarium sheets with forceps, and were softened in a solution of sulfosuccinate for three days (Erbar, 1995). The medial portion of the leaf blade has been shown to display the most regular arrangement of stomata (especially on the abaxial side (Rowson, 1946; Sharma and Dunn, 1968, 1969). Thus, the medial portion of each leaf blade was dissected out, sonicated for 2,5h in xylene to remove cuticular waxes, critical point dried, mounted on scanning electron microscope stubs, and sputter-coated with gold. Imaging was carried out on a JEOL JSM 6390 SEM at 10 kV.

Stomata and epidermis cells were counted manually on a leaf surface of 0.0775 mm^2^.

### 2.3. Data analysis

Tukey’s Honest Significance Difference test and Analyses of Variance (ANOVA) were implemented in R using the “MASS” and “car” packages on the SI and SD datasets to understand the change of these variables between Occident and Orient pre- and post- 1950.

## 3. Results

Table 1 and figure 2 summarise the results of the analyses of leaf traits (SI and SD) detailed in Supplementary file S1. Significant differences between pre- and post-1950 values of SI were identified for both leaf sides for both regions (Occident *versus* Orient), except for the occidental species on the adaxial leaf side. The stomata index significantly decreased in the occidental species (significant only for abaxial side), whereas a significant increase was observed in the oriental species (significant for both sides; see table 1 and figure 2a, b). Significant differences between pre- and post-1950 were not identified for changes in SD, except for oriental species on the abaxial side of the leaves (see table 1 and figure 2c, d).

**Figure 2.**
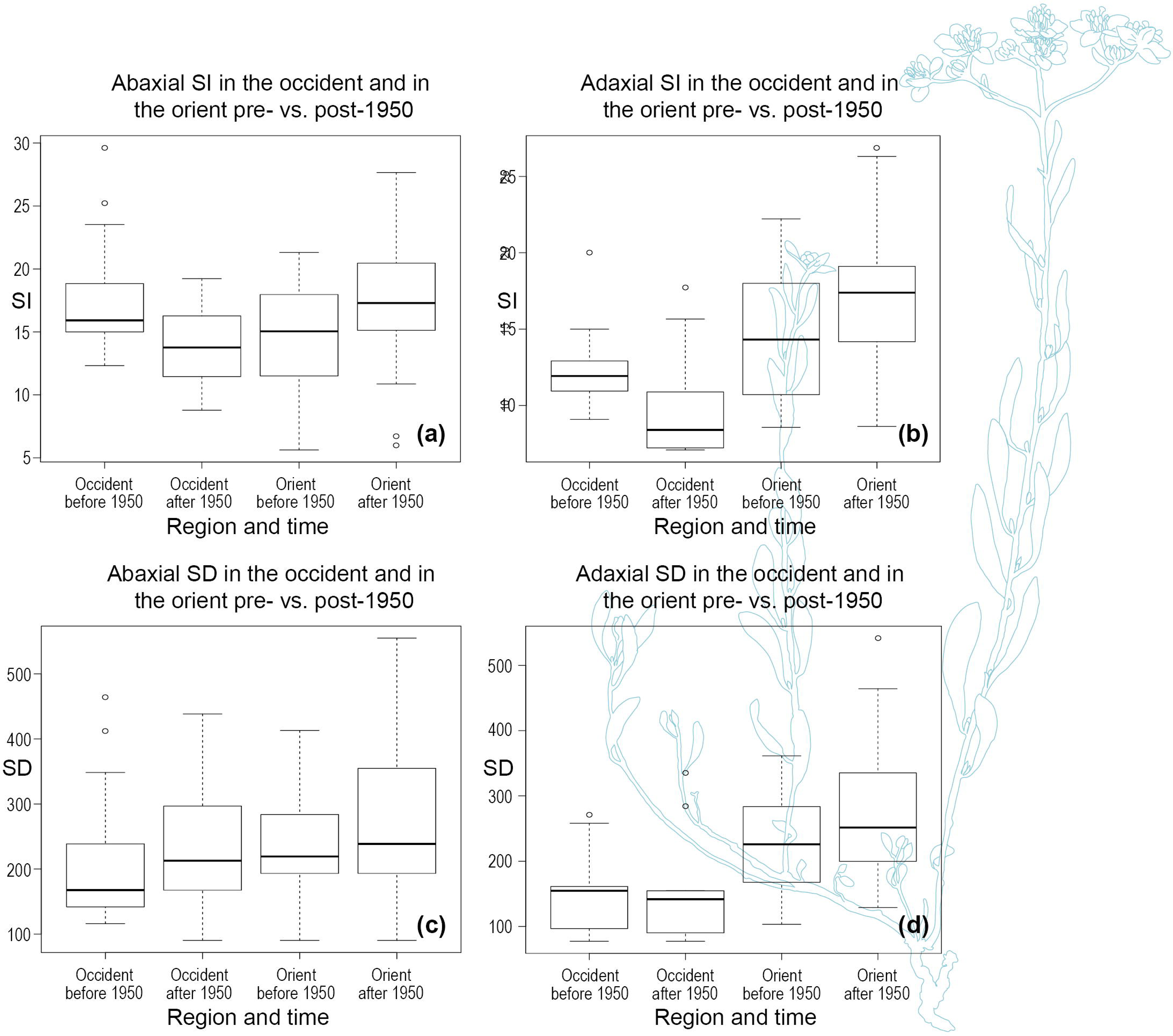
Stomata of *Haplophylum*. Boxplots of values of SI and SD before and after 1950, in Orient and in Occident. (a) SI values on the abaxial side of leaves. (b) SI values on the adaxial side of leaves. (c) SD values on the abaxial side of leaves. (d) SD values on the adaxial side of leaves. Background figure is *Haplophyllum villosum*, modified from (Townsend, 1986).

**Table 1.** Summary of results of ANOVA on significance and sign of differences between SI and SD of both leaf sides before and after 1950, for Occident and Orient.

| **Stomata trait** | **Difference** | **P value** | **Significance** |
| --- | --- | --- | --- |
| **Abaxial Stomata Index** |  |  |  |
| Occident after 1950 - Occident before 1950 | -4,6 | 0,003 | ** |
| Orient after 1950 - Orient before 1950 | 2,9 | 0,003 | ** |
| **Adaxial Stomata Index** |  |  |  |
| Occident after 1950 - Occident before 1950 | -1,7 | 0,437 |  |
| Orient after 1950 - Orient before 1950 | 2,4 | 0,007 | ** |
| **Abaxial Stomata Density** |  |  |  |
| Occident after 1950 - Occident before 1950 | -9,7 | 0,985 |  |
| Orient after 1950 - Orient before 1950 | 51,1 | 0,028 | * |
| **Adaxial Stomata Density** |  |  |  |
| Occident after 1950 - Occident before 1950 | 3,6 | 0,999 |  |
| Orient after 1950 - Orient before 1950 | 38,1 | 0,07 | . |

## 4. Discussion

Plants respond to the environmental signals they perceive by changes in their phenotype to increase individual fitness (Trewavas, 2009). The most conspicuous post-industrial environmental change that induces adaptations in plant phenotypes is the global increase in atmospheric CO_2_ concentration (de Boer et al., 2011; Ellis, 2015; Xu et al., 2015). Our results indicate that SI decreased between pre- and post-1950 in the occidental *Haplophyllum* species (significant only for abaxial side; table 1, Fig. 2a, b), but significantly increased between pre- and post-1950 in the oriental *Haplophyllum* species on both sides of the leaves (table a, Fig. 2a, b). Values for SD exhibited no significant trend possibly due to adaptation to other environmental conditions and variation in leaf expansion (table 1, Fig. 2c, d; (Royer, 2001)).

The decrease in SI observed in occidental species of *Haplophyllum* corroborates the studies on herbarium and fossil material over the past decades that overwhelmingly indicate an inverse relationship of SI with atmospheric CO_2_ concentration (e.g., Beerling and Chaloner, 1993; Royer, 2001). The increase in SI observed in oriental species of *Haplophyllum*, however, does not fit into this pattern. Other environmental factors must be responsible for this observation. Except CO_2_, other environmental factors that have been shown to influence stomata initiation are: irradiance (Furukawa, 1998; Gay and Hurd, 1975; Schoch et al., 1980; Sharma and Dunn, 1968, 1969), temperature (Ferris et al., 1996; Salisbury, 1928; Salisbury, 1927; Wagner, 1998), and drought stress (Makbul et al., 2011; Salisbury, 1927). Increases in all abovementioned factors have been shown to increase SI. These factors are expected to affect the oriental and occidental species of *Haplophyllum* differently because they grow in different climates and habitats: the occidental species grow in the mild Mediterranean climate under tree cover, whereas the oriental species of *Haplophyllum* grow in the harsh IT climate, which is characterised by strong continentality (Djamali et al., 2012), and a scarcity of trees (Takhtajan, 1986).

Total solar irradiance has increased during the period 1800-present (Solanki et al., 2013); this increase is expected to especially affect cloudless arid regions. Our results support a stronger influence of irradiance on oriental species: the oriental species display notably higher SI values for the sun-exposed, adaxial side of their leaves compared to their Mediterranean counterparts (Fig. 2b), whereas on the shaded, abaxial side of the leaves no such differences can be found (Fig. 2a). This notwithstanding, evidence of the positive effect of irradiance on SI was based on depletion experiments where the plants were exposed to a fraction of normal sunlight where a decrease of 94% and 99% in irradiance caused a 1,6x (Furukawa, 1998) and 1,5x (Sharma and Dunn, 1969) reduction in SI, respectively. Given that (1) the relationship between percent of total sunlight and SI is linear (Furukawa, 1998), and (2) the average increases (pre to post 1950) in SI of oriental species of *Haplophyllum* are 1,14x for abaxial side and 1,22x for adaxial side (table S1), our observations would require increases in total sun irradiance of 20% and 45% respectively. These differences in irradiance are two orders of magnitude higher than the percentual increase in irradiance reconstructed by the least conservative model reviewed by (Solanki et al., 2013), which proposes a 0,4% increase in irradiance since 1800 (Shapiro et al., 2011). Therefore, increases in irradiance do not appear to underlie the observed increase in SI since pre-industrial times, although experimental confirmation would be needed to test this.

Non-succulent xerophytes, such as *Haplophyllum* have been shown to possess increased SI with small stomata as an adaptation to rapidly changing water availability (Franks et al., 2009) and possibly also as an adaptation for the increased need for leaf cooling (Porporato et al., 2001). Therefore, increase in aridity, drought stress -a type of extreme events, whose frequency influence plants more than mean changes (Reyer et al., 2013)- and temperature, such as occurred in the Occident and Orient post-industrial revolution ((Hartmann et al., 2013), p208-222), would be expected to increase SI further. However, oriental (IT) and occidental (Mediterranean) plants do not grow under the same constraints. In the harsh, continental conditions that the oriental *Haplophyllum* species experience, plant growth models derived from long-term climate statistics (Nemani et al., 2003) predict that plant growth is mostly water and secondarily temperature limited, whereas the same models predict that the growth of their occidental counterparts are predominantly sunlight limited (Allen et al., 2010; Boisvenue and Running, 2006; Nemani et al., 2003; Running et al., 2004). Therefore the post-industrial increases in aridity, drought stress, and temperature that occurred both in the Mediterranean and IT regions ((Hartmann et al., 2013), p208-222) should have a stronger impact on the SI of plants whose growth is already water- and temperature-limited. The environment may thus constrain SI more tightly than atmospheric CO_2_, and paradoxically, a decrease in SI allowed by higher atmospheric CO_2_ would probably not be advantageous to the plants due to constraints of water stress response and leaf cooling.

In conclusion, we suggest that whereas atmospheric CO_2_ directly caused a decrease in SI in occidental xerophytes, it indirectly caused an increase in SI in oriental xerophytes *via* climate change (increase in aridity, drought stress, and temperature).

Potential evapotranspiration will increase with rising temperatures, and this will probably make the world effectively drier in many areas, independently of changes in precipitations (Cavender-Bares et al., 2016). Moreover, climate models project that arid and semiarid midlatitudes will become even drier than at present (Collins et al., 2013). Thus, in the future water stress will likely increase for all Eurasian xerophytes. Additional studies of the interaction between increased aridity and elevated CO_2_ are needed to enable us to better understand and predict not only the responses to climate change of the xerophytes from the Mediterranean and IT regions, but of global dryland as well (Maestre et al., 2016).

Finally, this study also highlights the value of historical collections to answer ecological questions, especially regarding climatic change. By drawing on collections from several herbaria, made over two centuries, we show how these data may provide valuable information even of poorly studied areas (e.g., IT floristic region), and we highlight the increasing value of natural history collections in understanding long-term changes (Pyke and Ehrlich, 2010).

## 5. Acknowledgments

The authors would like to thank Dr. Ernst Vitek (W) for making material ready for this study. Susanne Sontag and Andrea Frosch-Radivo are thanked for help in sample preparation and mounting. This research did not receive any specific grant from funding agencies in the public, commercial, or not-for-profit sectors.

---

**Table S1.**
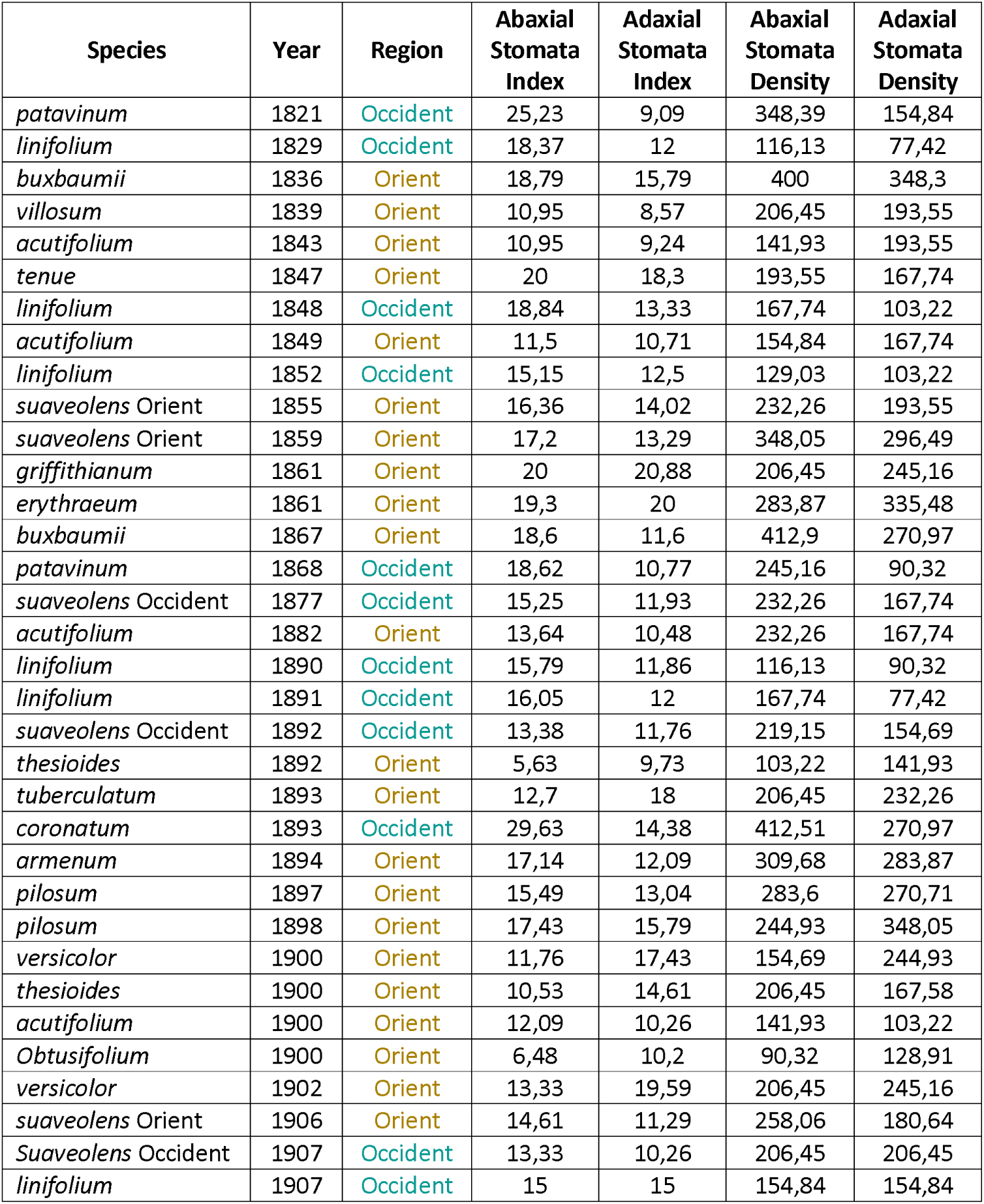
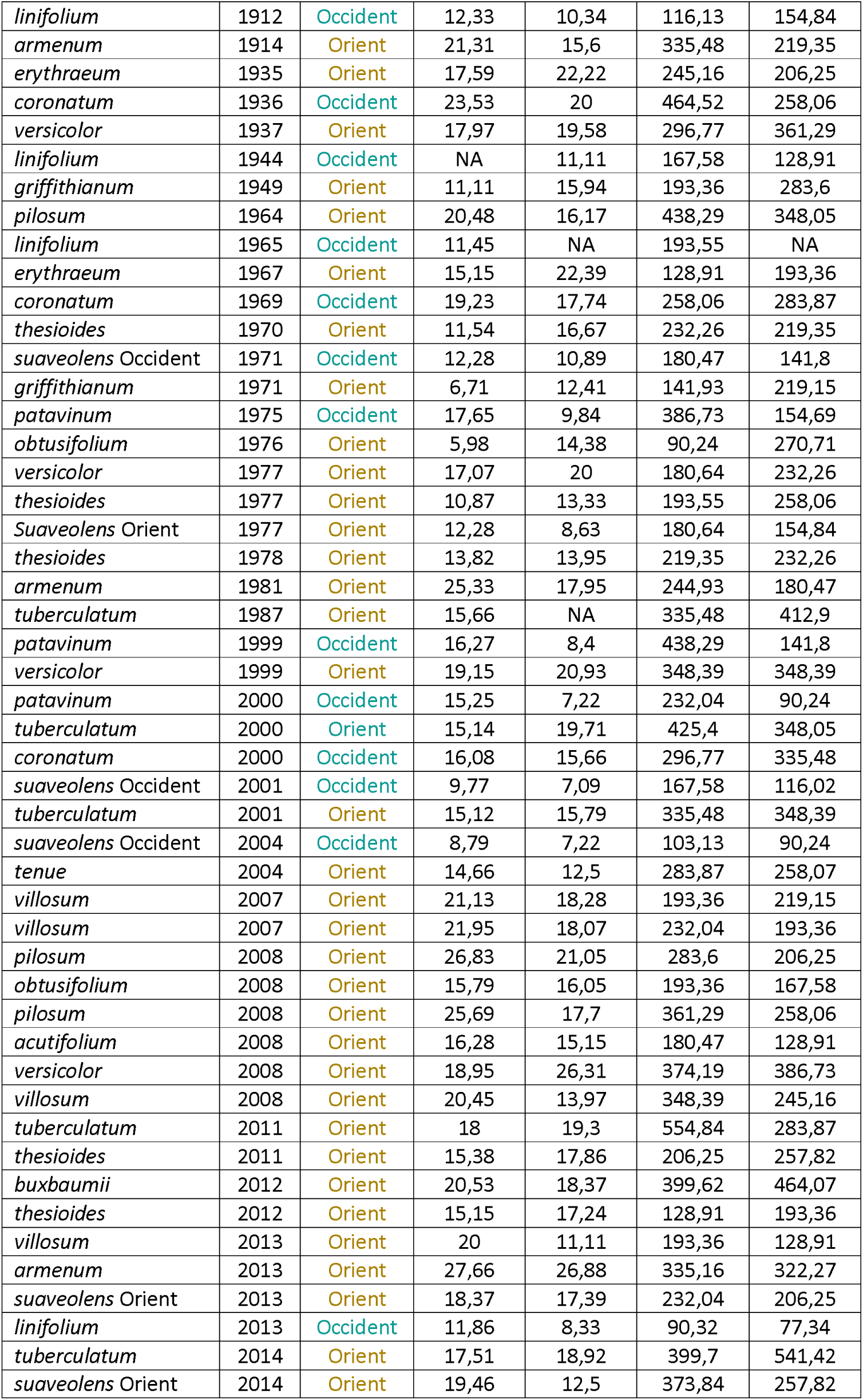
Dataset from 83 herbarium specimen of *Haplophyllum*. Stomata index and stomata density data from both leaf side, for 16 species, classified by region (Orient and Occident), and collection date (1821-2014). SI is obtained by the number of stomata per unit of leaf area divided to the number of epidermal cells plus number of stomata per unit leaf area; SD is obtained on a leaf surface of 0.0775 mm^2^.

## 6. References

Allen, C.D., Macalady, A.K., Chenchouni, H., Bachelet, D., McDowell, N., Vennetier, M., Kitzberger, T., Rigling, A., Breshears, D.D., Hogg, E.T., 2010. A global overview of drought and heat-induced tree mortality reveals emerging climate change risks for forests. Forest ecology and management 259, 660–684.

Bartholomeus, R.P., Witte, J.P.M., Runhaar, J., 2012. Drought stress and vegetation characteristics on sites with different slopes and orientations. Ecohydrology 5, 808–818.

Bates, B.C., Kundzewicz, Z.W., Wu, S., Palutikof, J.P., Eds., 2008. Climate change and water, Technical Paper of the Intergovernmental Panel on Climate Change. IPCC Secretariat, Geneva, p. 210.

Beerling, D.J., Chaloner, W.G., 1993. The Impact of Atmospheric Co2 and Temperature-Change on Stomatal Density - Observations from Quercus-Robur Lammas Leaves. Ann Bot-London 71, 231–235.

Beerling, D.J., Royer, D.L., 2002. Reading a CO2 signal from fossil stomata. New Phytologist 153, 387–397.

Boden, T., Andres, B., 2017. National CO2 Emissions from Fossil-Fuel Burning, Cement Manufacture, and Gas Flaring: 1751-2014, Carbon Dioxide Information Analysis Center, Oak Ridge National Laboratory. Appalachian State University, Oak Ridge, Tennessee, USA.

Boisvenue, C., Running, S.W., 2006. Impacts of climate change on natural forest productivity–evidence since the middle of the 20th century. Global Change Biology 12, 862–882.

Cavender-Bares, J., Ackerly, D.D., Hobbie, S.E., Townsend, P.A., 2016. Evolutionary Legacy Effects on Ecosystems: Biogeographic Origins, Plant Traits, and Implications for Management in the Era of Global Change. Annual Review of Ecology, Evolution, and Systematics 47, 433–462.

Collins, M., Knutti, R., Arblaster, J., Dufresne, J.-L., Fichefet, T., Friedlingstein, P., Gao, X., Gutowski, W.J., Johns, T., Krinner, G., Shongwe, M., Tebaldi, C., Weaver, A.J., Wehner, M., 2013. Chapter 12 - Long-term climate change: Projections, commitments and irreversibility, in: Ipcc (Ed.), Climate Change 2013: The Physical Science Basis. IPCC Working Group I Contribution to AR5. Cambridge University Press, Cambridge.

Crawley, T.J., Berner, R.A., 2001. CO2 and climate change. Science 292, 870–872.

de Boer, H.J., Lammertsma, E.I., Wagner-Cremer, F., Dilcher, D.L., Wassen, M.J., Dekker, S.C., 2011. Climate forcing due to optimization of maximal leaf conductance in subtropical vegetation under rising CO2. P Natl Acad Sci USA 108, 4041–4046.

Djamali, M., Brewer, S., Breckle, S.W., Jackson, S.T., 2012. Climatic determinism in phytogeographic regionalization: A test from the Irano-Turanian region, SW and Central Asia. Flora 207, 237–249.

Ellis, E.C., 2015. Ecology in an anthropogenic biosphere. Ecological Monographs 85, 287–331.

Erbar, C., 1995. On the floral development of Sphenoclea zeylanica (Sphenocleaceae, Campanulales): SEM investigations on herbarium material. Botanische Jahrbücher fur Systematik, Pflanzengeschichte und Pflanzengeographie 117, 469–483.

Ferris, R., Nijs, I., Behaeghe, T., Impens, I., 1996. Elevated CO 2 and temperature have different effects on leaf anatomy of perennial ryegrass in spring and summer. Ann Bot-London 78, 489–497.

Franks, P.J., Drake, P.L., Beerling, D.J., 2009. Plasticity in maximum stomatal conductance constrained by negative correlation between stomatal size and density: an analysis using Eucalyptus globulus. Plant Cell Environ 32, 1737–1748.

Furukawa, A., 1998. Stomatal frequency of Quercus myrsinaefolia grown under different irradiances. Photosynthetica 34, 195–199.

Gay, A., Hurd, R., 1975. The influence of light on stomatal density in the tomato. New Phytol 75, 37–46.

Greenwood, D.R., Scarr, M.J., Christophel, D.C., 2003. Leaf stomatal frequency in the Australian tropical rainforest tree Neolitsea dealbata (Lauraceae) as a proxy measure of atmospheric pCO(2). Palaeogeogr Palaeocl 196, 375–393.

Hartmann, D.L., Klein Tank, A.M.G., Rusticucci, M., Alexander, L.V., Brönnimann, S., Charabi, Y., Dentener, F.J., Dlugokencky, E.J., Easterling, D.R., Kaplan, A., Soden, B.J., Thorne, P.W., Wild, M., Zhai, P.M., 2013. Observations: Atmosphere and Surface, in: Stocker, T.F., Qin, D., Plattner, G.-K., Tignor, M., Allen, S.K., Boschung, J., Nauels, A., Xia, Y., Bex, V., Midgley, P.M. (Eds.), Climate Change 2013: The Physical Science Basis. Contribution of Working Group I to the Fifth Assessment Report of the Intergovernmental Panel on Climate Change Cambridge University Press, Cambridge, United Kingdom and New York, NY, USA.

Hibbard, K.A., Crutzen, P.J., Lambin, E.F., Liverman, D., Mantua, N.J., McNeill, J.R., Messerli, B., Steffen, W., 2006. Decadal interactions of humans and the environment, Integrated history and future of people on earth. Dahlem Workshop Report, pp. 341–375.

Kreft, H., Jetz, W., 2010. A framework for delineating biogeographical regions based on species distributions. J Biogeogr 37, 2029–2053.

Kürschner, W.M., Wagner, F., Dilcher, D.L., Visscher, H., 2001. Using fossil leaves for the reconstruction of Cenozoic paleoatmospheric CO2 concentration, in: Gerhard, L.C., Harrison, W.E., Hanson, B.M. (Eds.), Geological Perspectives of Global Climate Change. Am. Assoc. Pet. Geol., Boulder, OK, USA, pp 155–176.

Maestre, F.T., Eldridge, D.J., Soliveres, S., Kéfi, S., Delgado-Baquerizo, M., Bowker, M.A., García-Palacios, P., Gaitán, J., Gallardo, A., Lázaro, R., 2016. Structure and functioning of dryland ecosystems in a changing world. Annual review of ecology, evolution, and systematics 47, 215–237.

Makbul, S., GÜLER, N.S., DURMUŞ, N., GÜVEN, S., 2011. Changes in anatomical and physiological parameters of soybean under drought stress. Turkish Journal of Botany 35, 369–377.

Manafzadeh, S., Salvo, G., Conti, E., 2014. A tale of migrations from east to west: the Irano-Turanian floristic region as a source of Mediterranean xerophytes. J Biogeogr 41, 366–379.

Manafzadeh, S., Staedler, Y.M., Conti, E., 2016. Visions of the past and dreams of the future in the Orient: the Irano-Turanian region from classical botany to evolutionary studies. Biol Rev.

Maximov, N.A., 1929. The plant in relation to water. George Allen and Unwin Ltd, London, UK.

McNeill, J.R., Engelke, P., 2016. The great acceleration. Harvard University Press.

Nemani, R.R., Keeling, C.D., Hashimoto, H., Jolly, W.M., Piper, S.C., Tucker, C.J., Myneni, R.B., Running, S.W., 2003. Climate-driven increases in global terrestrial net primary production from 1982 to 1999. Science 300, 1560–1563.

Pachauri, R.K., Allen, M.R., Barros, V.R., Broome, J., Cramer, W., Christ, R., Church, J.A., Clarke, L., Dahe, Q., Dasgupta, P., 2014. Climate change 2014: synthesis report. Contribution of Working Groups I, II and III to the fifth assessment report of the Intergovernmental Panel on Climate Change. IPCC.

Porporato, A., Laio, F., Ridolfi, L., Rodriguez-Iturbe, I., 2001. Plants in water-controlled ecosystems: active role in hydrologic processes and response to water stress - III. Vegetation water stress. Adv Water Resour 24, 725–744.

Pyke, G.H., Ehrlich, P.R., 2010. Biological collections and ecological/environmental research: a review, some observations and a look to the future. Biol Rev 85, 247–266.

Retallack, G.J., 2001. A 300-million-year record of atmospheric carbon dioxide from fossil plant cuticles. Nature 411, 287–290.

Reyer, C.P., Leuzinger, S., Rammig, A., Wolf, A., Bartholomeus, R.P., Bonfante, A., de Lorenzi, F., Dury, M., Gloning, P., Abou Jaoudé, R., 2013. A plant’s perspective of extremes: terrestrial plant responses to changing climatic variability. Global change biology 19, 75–89.

Rowson, J.M., 1946. The significance of stomatal index as a differential character. Part III. Studies in the genera Atropa, Datura, Digitalis, Phytolacca and in polyploid leaves. Quart. J. Pharm. Pharmaceut. 19, 136–143.

Royer, D.L., 2001. Stomatal density and stomatal index as indicators of paleoatmospheric CO2 concentration. Rev Palaeobot Palyno 114, 1–28.

Running, S.W., Nemani, R.R., Heinsch, F.A., Zhao, M., Reeves, M., Hashimoto, H., 2004. A continuous satellite-derived measure of global terrestrial primary production. AIBS Bulletin 54, 547–560.

Salisbury, E., 1928. On the causes and ecological significance of stomatal frequency, with special reference to the woodland flora. Philosophical Transactions of the Royal Society of London. Series B, Containing Papers of a Biological Character 216, 1–65.

Salisbury, E.J., 1927. On the causes and ecological significance of stomatal frequency, with special reference to the woodland flora. Philosophical Transactions of the Royal Society London B 216, 1–65.

Salvo, G., Manafzadeh, S., Ghahremaninejad, F., Tojibaev, K., Zeltner, L., Conti, E., 2011. Phylogeny, morphology, and biogeography of Haplophyllum (Rutaceae), a species-rich genus of the Irano-Turanian floristic region. Taxon 60, 513–527.

Schoch, P.-G., Zinsou, C., Sibi, M., 1980. Dependence of the stomatal index on environmental factors during stomatal differentiation in leaves of Vigna sinensis L. 1. Effect of light intensity. Journal of Experimental Botany 31, 1211–1216.

Shapiro, A., Schmutz, W., Rozanov, E., Schoell, M., Haberreiter, M., Shapiro, A., Nyeki, S., 2011. A new approach to the long-term reconstruction of the solar irradiance leads to large historical solar forcing. Astronomy & Astrophysics 529, A67.

Sharma, G.K., Dunn, D.B., 1968. Effect of Environment on Cuticular Features in Kalanchoe Fedschenkoi. B Torrey Bot Club 95, 464-&.

Sharma, G.K., Dunn, D.B., 1969. Environmental Modifications of Leaf Surface Traits in Datura Stramonium. Can J Botany 47, 1211-&.

Solanki, S.K., Krivova, N.A., Haigh, J.D., 2013. Solar irradiance variability and climate. Annual Review of Astronomy and Astrophysics 51, 311–351.

Solomon, S., Qin, D., Manning, M., Alley, R.B., Berntsen, T., Bindoff, N.L., Chen, Z., Chidthaisong, A., Gregory, J.M., Hegerl, G.C., Heimann, M., Hewitson, B., Hoskins, B.J., Joos, F., Jouzel, J., Kattsov, V., Lohmann, U., Matsuno, T., Molina, M., Nicholls, N., Overpeck, J., Raga, G., Ramaswamy, V., Ren, J., Rusticucci, M., Somerville, R., Stocker, T.F., Whetton, P., Wood, R., Wratt, D., 2007. Technical summary, in: Solomon S, Q.D., Manning M, et al. (Ed.), Climate Change 2007: The Physical Science Basis. Contribution of Working Group I to the Fourth Assessment Report of the Intergovernmental Panel on Climate Change. Cambridge University Press, Cambridge, United Kingdom and New York, NY, USA.

Steffen, W., Broadgate, W., Deutsch, L., Gaffney, O., Ludwig, C., 2015. The trajectory of the Anthropocene: the great acceleration. The Anthropocene Review 2, 81–98.

Takhtajan, A., 1986. Floristic regions of the world. University of California Press, Berkeley, CA, USA.

Townsend, C.C., 1986. Taxonomic revision of the genus Haplophyllum (Rutaceae). Bentham-Moxon, Kent U.K.

Trewavas, A., 2009. What is plant behaviour? Plant Cell Environ 32, 606–616.

van der Burgh, J., Visscher, H., Dilcher, D.L., Kürschner, W.M., 1993. Paleoatmosphere signatures in Neogene fossil leaves. Science 260, 1788–1790.

Volkens, G., 1887. Die Flora der Aegyptisch-arabischen Wüste auf Grundlage anatomischphysiologischer Forschungen dargestellt. Gebruder Borntraeger, Berlin, Germany.

Wagner, F., 1998. The influence of environment on the stomatal frequency in Betula. LPP Contrib. Series 9, 102.

Wegehenkel, M., 2009. Modeling of vegetation dynamics in hydrological models for the assessment of the effects of climate change on evapotranspiration and groundwater recharge. Advances in Geosciences 21, 109–115.

Willmer, C., Fricker, M., 1996. Stomata, 2nd edition ed. Chapman & Hall, London, UK.

Xu, Z., Jiang, Y., Zhou, G., 2015. Response and adaptation of photosynthesis, respiration, and antioxidant systems to elevated CO2 with environmental stress in plants. Frontiers in plant science 6.

Zohary, M., 1973. Geobotanical foundations of the Middle East. Gustav Fischer, Stuttgart, Germany.

