## Supplementary Materials for "Response of oriental xerophytes to the occidental industrial revolution"

```
########################  1  ############################
Analysis of Variance Table

Response: Abaxial Stomata Index
                    Df  Sum Sq Mean Sq F value    Pr(>F)    
recentPeriod         1   11.98  11.977  1.2783    0.2625    
Region               1    0.17   0.175  0.0186    0.8919    
Species             15 1016.13  67.742  7.2301 6.978e-09 ***
recentPeriod:Region  1  264.00 264.005 28.1775 1.529e-06 ***
Residuals           63  590.27   9.369                      
---
Signif. codes:  0 ‘***’ 0.001 ‘**’ 0.01 ‘*’ 0.05 ‘.’ 0.1 ‘ ’ 1
-------------------------- lm -------------------------------

Call:
lm(formula = `Abaxial Stomata Index` ~ recentPeriod * Region + 
    Species, data = dat)

Residuals:
    Min      1Q  Median      3Q     Max 
-8.0225 -1.2373 -0.0313  1.4607  8.4562 

Coefficients:
                               Estimate Std. Error t value Pr(>|t|)    
(Intercept)                   12.245003   2.347687   5.216 2.17e-06 ***
recentPeriodTRUE              -5.286110   1.335198  -3.959 0.000194 ***
RegionOrient                   0.009254   1.939163   0.005 0.996207    
Speciesarmenum                 9.011386   2.070065   4.353 5.02e-05 ***
Speciesbuxbaumii               5.989505   2.238440   2.676 0.009491 ** 
Speciescoronatum              12.515552   2.721817   4.598 2.11e-05 ***
Specieserythraeum              4.029505   2.238440   1.800 0.076626 .  
Speciesgriffithianum          -0.710495   2.238440  -0.317 0.751985    
Specieslinifolium              3.911911   2.498909   1.565 0.122488    
SpeciesObtusifolium           -4.963400   2.272410  -2.184 0.032677 *  
Speciespatavinum               9.530663   2.637735   3.613 0.000601 ***
Speciespilosum                 7.016514   1.967316   3.567 0.000697 ***
Speciessuaveolens              2.531386   1.872001   1.352 0.181134    
Speciestenue                   3.481386   2.574394   1.352 0.181110    
Speciesthesioides             -2.686196   1.847972  -1.454 0.151023    
Speciestuberculatum            0.776814   1.934621   0.402 0.689387    
Speciesversicolor              2.523052   1.872001   1.348 0.182555    
Speciesvillosum                4.090771   2.005884   2.039 0.045611 *  
recentPeriodTRUE:RegionOrient  8.474825   1.596540   5.308 1.53e-06 ***
---
Signif. codes:  0 ‘***’ 0.001 ‘**’ 0.01 ‘*’ 0.05 ‘.’ 0.1 ‘ ’ 1

Residual standard error: 3.061 on 63 degrees of freedom
  (1 observation deleted due to missingness)
Multiple R-squared:  0.6865,	Adjusted R-squared:  0.5969 
F-statistic: 7.663 on 18 and 63 DF,  p-value: 5.189e-10

-------------------------- * ------------------------------
                                  diff        lwr        upr       p adj
TRUE:Occident-FALSE:Occident -4.5642785 -7.9087543 -1.2198026 0.003412187
FALSE:Orient-FALSE:Occident  -3.5254349 -6.2031587 -0.8477112 0.005034993
TRUE:Orient-FALSE:Occident   -0.5777996 -3.1661697  2.0105705 0.935021697
FALSE:Orient-TRUE:Occident    1.0388436 -1.9668939  4.0445810 0.798481856
TRUE:Orient-TRUE:Occident     3.9864788  1.0600624  6.9128952 0.003482124
TRUE:Orient-FALSE:Orient      2.9476353  0.8148909  5.0803796 0.002958749
########################  2  ############################
Analysis of Variance Table

Response: Adaxial Stomata Index
                    Df Sum Sq Mean Sq F value    Pr(>F)    
recentPeriod         1  66.59  66.588  9.1461  0.003623 ** 
Region               1 281.18 281.182 38.6212 4.813e-08 ***
Species             15 666.02  44.401  6.0987 1.375e-07 ***
```

```
recentPeriod:Region  1  82.66  82.665 11.3543  0.001299 **
Residuals           62 451.39   7.280
---
Signif. codes:  0 ‘***’ 0.001 ‘**’ 0.01 ‘*’ 0.05 ‘.’ 0.1 ‘ ’ 1
-------------------------- lm -------------------------------

Call:
lm(formula = `Adaxial Stomata Index` ~ recentPeriod * Region +
    Species, data = dat)

Residuals:
    Min      1Q  Median      3Q     Max
-5.8526 -1.5442 -0.0745  1.6096  7.3606

Coefficients:
                              Estimate Std. Error t value Pr(>|t|)
(Intercept)                    10.1068     2.0817   4.855 8.50e-06 ***
recentPeriodTRUE               -2.2003     1.2596  -1.747  0.08561 .
RegionOrient                    0.5054     1.7242   0.293  0.77041
Speciesarmenum                  6.1283     1.8248   3.358  0.00134 **
Speciesbuxbaumii                3.7148     1.9732   1.883  0.06444 .
Speciescoronatum                7.9383     2.3993   3.309  0.00156 **
Specieserythraeum               9.9981     1.9732   5.067 3.90e-06 ***
Speciesgriffithianum            4.8715     1.9732   2.469  0.01633 *
Specieslinifolium               1.9677     2.2328   0.881  0.38159
SpeciesObtusifolium             1.0785     2.0032   0.538  0.59224
Speciespatavinum                0.2774     2.3256   0.119  0.90545
Speciespilosum                  4.4704     1.7343   2.578  0.01234 *
Speciessuaveolens               0.8517     1.6502   0.516  0.60763
Speciestenue                    3.3983     2.2694   1.497  0.13934
Speciesthesioides               2.1729     1.6291   1.334  0.18717
Speciestuberculatum             5.5087     1.7684   3.115  0.00278 **
Speciesversicolor               8.6383     1.6502   5.235 2.08e-06 ***
Speciesvillosum                 1.1647     1.7684   0.659  0.51259
recentPeriodTRUE:RegionOrient   4.9792     1.4777   3.370  0.00130 **
---
Signif. codes:  0 ‘***’ 0.001 ‘**’ 0.01 ‘*’ 0.05 ‘.’ 0.1 ‘ ’ 1

Residual standard error: 2.698 on 62 degrees of freedom
  (2 observations deleted due to missingness)
Multiple R-squared:  0.7084,    Adjusted R-squared:  0.6237
F-statistic: 8.367 on 18 and 62 DF,  p-value: 1.114e-10

-------------------------- * ------------------------------
                                  diff         lwr      upr        p adj
TRUE:Occident-FALSE:Occident -1.719642 -4.72322694 1.283943 4.368814e-01
FALSE:Orient-FALSE:Occident   2.384378  0.07465055 4.694106 4.049544e-02
TRUE:Orient-FALSE:Occident    4.790284  2.54973962 7.030829 2.598563e-06
FALSE:Orient-TRUE:Occident    4.104020  1.34898442 6.859056 1.200981e-03
TRUE:Orient-TRUE:Occident     6.509926  3.81262749 9.207225 1.542608e-07
TRUE:Orient-FALSE:Orient      2.405906  0.51150860 4.300303 7.288403e-03
########################  3  ############################
Analysis of Variance Table

Response: Abaxial Stomata Density
                    Df Sum Sq Mean Sq F value    Pr(>F)
recentPeriod         1  27739   27739  6.0969   0.01622 *
Region               1  12240   12240  2.6904   0.10586
Species             15 519101   34607  7.6065 2.454e-09 ***
recentPeriod:Region  1  17845   17845  3.9223   0.05195 .
Residuals           64 291176    4550
---
Signif. codes:  0 ‘***’ 0.001 ‘**’ 0.01 ‘*’ 0.05 ‘.’ 0.1 ‘ ’ 1
-------------------------- lm -------------------------------

Call:
lm(formula = `Abaxial Stomata Density` ~ recentPeriod * Region +
```

```
    Species, data = dat)

Residuals:
     Min       1Q   Median       3Q      Max 
-142.970  -43.832    1.392   36.231  173.254 

Coefficients:
                              Estimate Std. Error t value Pr(>|t|)    
(Intercept)                    112.613     51.716   2.178 0.033135 *  
recentPeriodTRUE               -37.305     29.299  -1.273 0.207537    
RegionOrient                    51.240     42.710   1.200 0.234677    
Speciesarmenum                 126.377     45.616   2.770 0.007319 ** 
Speciesbuxbaumii               229.598     49.326   4.655 1.68e-05 ***
Speciescoronatum               264.004     59.978   4.402 4.16e-05 ***
Specieserythraeum               44.738     49.326   0.907 0.367814    
Speciesgriffithianum             6.005     49.326   0.122 0.903484    
Specieslinifolium               36.767     54.701   0.672 0.503912    
SpeciesObtusifolium            -60.657     50.075  -1.211 0.230224    
Speciespatavinum               239.892     58.125   4.127 0.000108 ***
Speciespilosum                 139.189     43.352   3.211 0.002072 ** 
Speciessuaveolens               90.879     41.251   2.203 0.031198 *  
Speciestenue                    58.774     56.729   1.036 0.304081    
Speciesthesioides               -2.544     40.722  -0.062 0.950372    
Speciestuberculatum            185.567     42.631   4.353 4.94e-05 ***
Speciesversicolor               80.252     41.251   1.945 0.056116 .  
Speciesvillosum                 45.134     44.202   1.021 0.311053    
recentPeriodTRUE:RegionOrient   69.471     35.078   1.980 0.051953 .  
---
Signif. codes:  0 ‘***’ 0.001 ‘**’ 0.01 ‘*’ 0.05 ‘.’ 0.1 ‘ ’ 1

Residual standard error: 67.45 on 64 degrees of freedom
Multiple R-squared:  0.6646,    Adjusted R-squared:  0.5702 
F-statistic: 7.045 on 18 and 64 DF,  p-value: 2.191e-09

-------------------------- * ------------------------------
                                   diff        lwr       upr      p adj
TRUE:Occident-FALSE:Occident -9.6611585 -82.298561  62.97624 0.98504337
FALSE:Orient-FALSE:Occident  -0.2738846 -57.963223  57.41545 0.99999929
TRUE:Orient-FALSE:Occident   50.7827538  -4.892761 106.45827 0.08613113
FALSE:Orient-TRUE:Occident    9.3872739 -56.819208  75.59376 0.98199812
TRUE:Orient-TRUE:Occident    60.4439123  -4.015388 124.90321 0.07386663
TRUE:Orient-FALSE:Orient     51.0566384   4.079315  98.03396 0.02798061
########################  4  ############################
Analysis of Variance Table

Response: Adaxial Stomata Density
                    Df Sum Sq Mean Sq F value    Pr(>F)    
recentPeriod         1  36309   36309 10.8847  0.001598 ** 
Region               1 145082  145082 43.4924 1.010e-08 ***
Species             15 311850   20790  6.2324 8.722e-08 ***
recentPeriod:Region  1   5835    5835  1.7492  0.190754    
Residuals           63 210156    3336                      
---
Signif. codes:  0 ‘***’ 0.001 ‘**’ 0.01 ‘*’ 0.05 ‘.’ 0.1 ‘ ’ 1
-------------------------- lm ------------------------------

Call:
lm(formula = `Adaxial Stomata Density` ~ recentPeriod * Region + 
    Species, data = dat)

Residuals:
     Min       1Q   Median       3Q      Max 
-106.356  -31.725   -4.863   38.588  175.765 

Coefficients:
                              Estimate Std. Error t value Pr(>|t|)    
(Intercept)                      98.96      44.56   2.221 0.029961 *  
```

```
recentPeriodTRUE                -14.78      26.96  -0.548 0.585537    
RegionOrient                     47.87      36.90   1.297 0.199341    
Speciesarmenum                   91.15      39.06   2.334 0.022826 *  
Speciesbuxbaumii                205.28      42.24   4.860 8.14e-06 ***
Speciescoronatum                195.53      51.36   3.807 0.000321 ***
Specieserythraeum                89.19      42.24   2.112 0.038682 *  
Speciesgriffithianum             93.47      42.24   2.213 0.030535 *  
Specieslinifolium                10.19      47.79   0.213 0.831897    
SpeciesObtusifolium              24.22      42.88   0.565 0.574233    
Speciespatavinum                 36.29      49.78   0.729 0.468737    
Speciespilosum                  123.18      37.12   3.318 0.001508 ** 
Speciessuaveolens                54.59      35.32   1.545 0.127255    
Speciestenue                     52.56      48.58   1.082 0.283358    
Speciesthesioides                43.91      34.87   1.259 0.212539    
Speciestuberculatum             191.79      36.50   5.254 1.88e-06 ***
Speciesversicolor               142.78      35.32   4.042 0.000147 ***
Speciesvillosum                  27.57      37.85   0.728 0.469046    
recentPeriodTRUE:RegionOrient    41.82      31.62   1.323 0.190754    
---
Signif. codes:  0 ‘***’ 0.001 ‘**’ 0.01 ‘*’ 0.05 ‘.’ 0.1 ‘ ’ 1

Residual standard error: 57.76 on 63 degrees of freedom
  (1 observation deleted due to missingness)
Multiple R-squared:  0.7037,    Adjusted R-squared:  0.619 
F-statistic: 8.312 on 18 and 63 DF,  p-value: 1.056e-10

-------------------------- * ------------------------------
                                  diff        lwr       upr        p adj
TRUE:Occident-FALSE:Occident   3.571688 -60.692734  67.83611 9.988630e-01
FALSE:Orient-FALSE:Occident   78.997328  29.578608 128.41605 4.568442e-04
TRUE:Orient-FALSE:Occident   117.102667  69.409059 164.79628 9.522153e-08
FALSE:Orient-TRUE:Occident    75.425640  16.479148 134.37213 6.746792e-03
TRUE:Orient-TRUE:Occident    113.530979  56.023073 171.03888 1.301702e-05
TRUE:Orient-FALSE:Orient      38.105339  -2.137092  78.34777 6.981643e-02
```
